# Origin and functional impact of early nonlinearities in primate retina

**DOI:** 10.64898/2026.03.19.713068

**Authors:** Vyom Raval, Rachel Oaks-Leaf, Qiang Chen, Fred Rieke

**Affiliations:** Department of Neurobiology and Biophysics Graduate Program in Neuroscience University of Washington, Seattle, WA 98195

## Abstract

Receptive fields provide a concise description of the stimulus selectivity of visual neurons. But this stimulus selectivity is neither static nor linear, and these nonlinear effects are not well captured by standard linear or pseudo-linear receptive field models. At the same time, receptive field models incorporating nonlinear effects are largely empirical, and are not easily interpreted in terms of underlying cellular and synaptic mechanisms. Here we show that two nonlinear mechanisms in the primate outer retina shape neural responses and that these contribute significantly to responses to natural stimuli and to the retinal output signals. Incorporating these outer retinal nonlinearities into models for visual function will improve our ability to identify the mechanistic origin of specific features of downstream visual responses.

## Introduction

Neural computation relies on specific combinations of linear and nonlinear processing steps. This is clear from classic computational models such as motion energy models (Adelson and Bergen, 1985), models for divisive normalization (Heeger et al., 1996), and winner-take-all models such as those used to study decision making (Shadlen and Kiani, 2013). Many of these models have been developed to account for specific computations and the correspondence of model components with biological mechanisms is unclear. A deeper understanding of the mechanistic basis of neural computation requires linking these empirical and specialized descriptions to the underlying biological mechanisms.

Retinal circuits exemplify this issue. One well-studied example comes from spatially localized nonlinearities that produce receptive field subunits (Enroth-Cugell and Robson, 1966; Hochstein and Shapley, 1976). Subunits confer sensitivity to spatial structure—allowing retinal ganglion cells to respond not just to the average light intensity in their receptive field, but to the spatial arrangement of light. The resulting nonlinear integration of spatial inputs shapes ganglion cell responses, including responses to motion (Demb et al., 2001b; Kuo et al., 2016; Manookin et al., 2018) and natural images (Turner and Rieke, 2016). Receptive field subunits also provide a key component of models for several specialized retinal computations such as differential motion detection (Baccus et al., 2008), latency coding (Gollisch and Meister, 2008) and predictive coding (Liu et al., 2021).

Two sequential processing layers shape retinal signaling. The outer retina consists of photoreceptors, horizontal cells and bipolar cell dendrites, while the inner retina consists of bipolar axon terminals, amacrine cells and ganglion cell dendrites. Receptive field subunits are often attributed to nonlinear synaptic processing between bipolar and ganglion cells. The evidence for this comes from experiments showing that nonlinear subunits are already present in a ganglion cell’s excitatory synaptic inputs (Demb et al., 2001a, 1999). In this view, the outer retina is treated as linear or near-linear, with key nonlinear steps confined to the inner retina.

Several findings challenge the assumption that the outer retina acts linearly. First, at low light levels, rod photoreceptor signals are thresholded before reaching the inner retina (Field and Rieke, 2002; Sampath and Rieke, 2004); this nonlinear processing in the outer retina is an essential component of vision in starlight. Second, photoreceptors exhibit adaptive temporal nonlinearities that shape downstream responses (Angueyra et al., 2022; Chen et al., 2024; Endeman and Kamermans, 2010; Yu et al., 2022). Third, dendritic inputs to bipolar cells in the salamander retina show nonlinear integration of spatial inputs, indicating that nonlinear subunits can arise prior to transmitter release at the bipolar output synapse (Schreyer and Gollisch, 2021). Nonetheless, we lack a clear understanding of the functional properties of nonlinear processing in the outer retina and how these properties contribute to retinal outputs, particularly in primate retina.

We show here that signals in the outer primate retina exhibit clear spatial and temporal nonlinearities that shape responses to naturalistic visual inputs. We find that at least two mechanisms contribute to this processing: dynamic nonlinearities in cone phototransduction and an additional nonlinear transformation at the cone output synapse. We develop models that capture these nonlinearities and accurately predict responses to a variety of stimuli. Finally, we provide several examples that illustrate the impact of these outer retina nonlinearities on responses of retinal ganglion cells.

## Results

We first identify and characterize two distinct mechanisms that contribute to nonlinear responses in horizontal and bipolar cells. We then examine how these mechanisms shape the encoding of complex stimuli, including naturalistic movies. Finally, we describe two ways in which these outer retina nonlinearities impact retinal ganglion cell responses. We focus on responses of horizontal cells since their synaptic inputs originate entirely in the outer retina and hence they provide a means to characterize outer retinal signals separately from inner retinal mechanisms.

### Nonlinearity at cone output synapse contributes to horizontal and bipolar responses

Figure 1A compares responses of cone photoreceptors, horizontal cells and bipolar cells to spatially-homogeneous Gaussian noise stimuli; all stimuli had the same mean light level and contrast. We fit these responses with linear-nonlinear models to determine whether nonlinearities in the stimulus-to-response transformation contributed substantially to the measured responses for these stimulus conditions.

**Figure 1:**
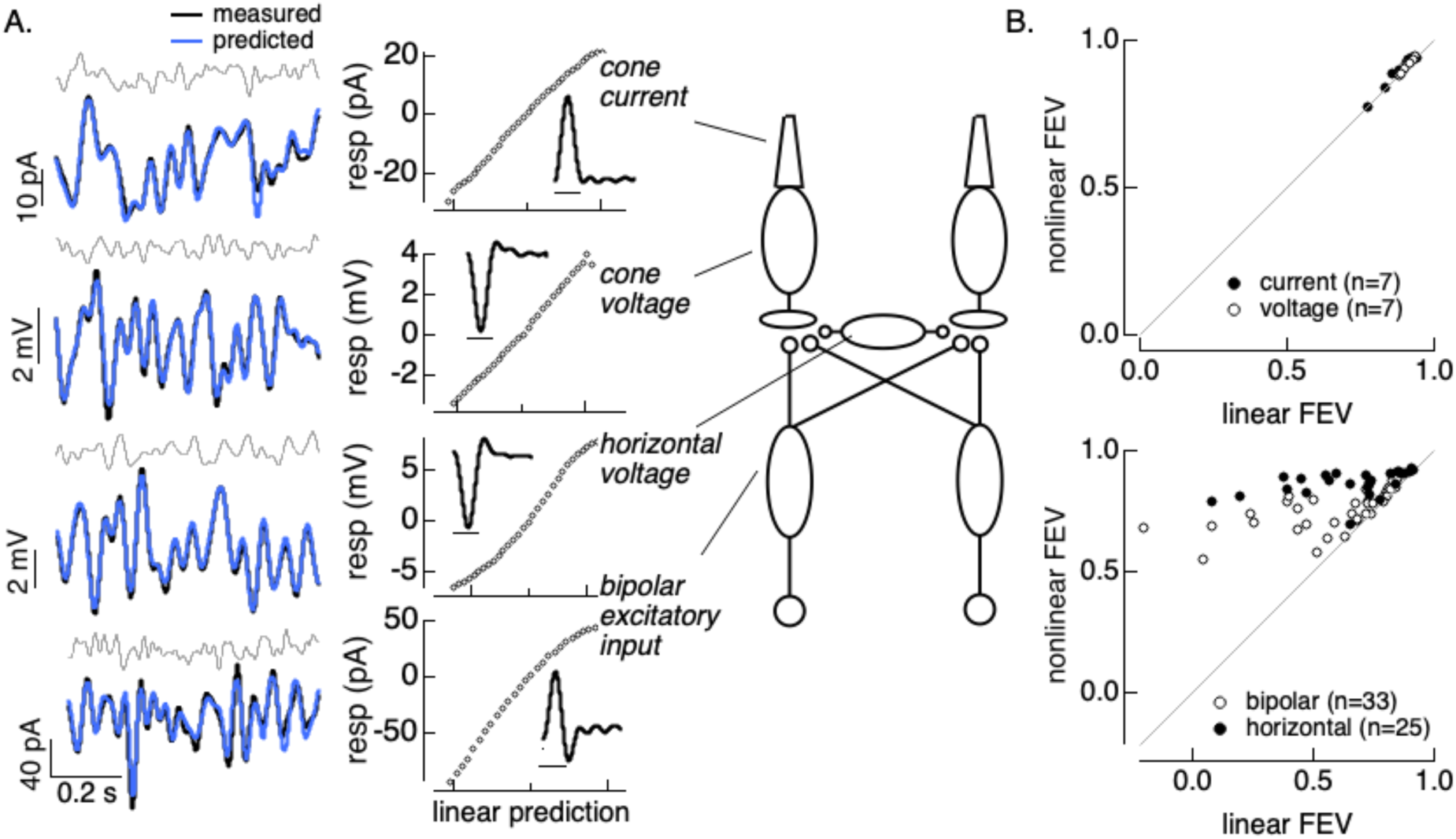
Post-transduction nonlinearity shapes horizontal and bipolar responses. A. Responses to gaussian noise stimuli and linear-nonlinear fits across neurons in the outer retina. B. Comparison of fraction explained variance for linear vs linear-nonlinear models for cones (top) and horizontal and bipolar responses (bottom). Horizontal responses are a combination of current and voltage measurements; bipolar responses are all excitatory synaptic currents. Stimuli all had a mean of 15,000 R*/cone/s and 50% contrast.

Cone photocurrents and voltages depended linearly or near-linearly on the stimuli, as indicated by the lack of prominent curvature in the estimated nonlinearities (Figure 1A, top two rows). We quantified potential nonlinearities on the stimulus-response relation by comparing the fraction of variance in the measured responses that was explained by linear and nonlinear models; adding a nonlinearity had little or no impact on the accuracy of predictions of cone photocurrents or photovoltages elicited by noise stimuli (Figure 1B, top). Thus cone photoreceptors respond linearly or near-linearly to Gaussian noise stimuli at a single mean light level (see also (Angueyra et al., 2022); we return below to higher contrast stimuli that reveal nonlinearities in cone responses (Angueyra et al., 2022; Chen et al., 2024; Endeman and Kamermans, 2010).

We compared cone responses to horizontal cell voltages measured under current clamp (Figure 1A, second from bottom) and bipolar cell excitatory synaptic inputs (Figure 1A, bottom) measured under voltage clamp (see Methods). Horizontal cell and bipolar cell responses exhibited clear nonlinearities for the same stimuli, as indicated by the curvature in the LN model nonlinearities (Figure 1A, bottom two rows). Consistent with these examples, including a nonlinearity systematically improved predictions of horizontal and bipolar responses (Figure 1B, bottom). Excitatory synaptic inputs to horizontal cells measured under voltage clamp showed a similar nonlinearity (not shown). The linearity of the cone photovoltages and nonlinearity in the bipolar and horizontal cell synaptic inputs indicates that transmission of signals across the cone output synapse is nonlinear.

### Nonlinearity in cone phototransduction contributes to horizontal and bipolar responses

Figure 1 evaluates responses at a single mean light level. To probe how responses changed with changes in light level, we compared linear-nonlinear models fit to horizontal and bipolar cell responses at three light levels. These spanned a 10-fold range of light levels, with the lowest near the onset of adaptation in cone phototransduction (Angueyra and Rieke, 2013; Cao et al., 2014; Schneeweis and Schnapf, 1999). Figure 2A shows linear filters and nonlinearities from LN models fit to horizontal cell responses at the three light levels. In addition to the curvature in the nonlinearities described above, the time course of the linear filter speeds and the slope of the nonlinearity decreases systematically with increases in mean light level. We quantified changes in response kinetics from the time-to-peak of the linear filter and changes in gain from the slope of the nonlinearity (Figure 2C and D, black lines).

**Figure 2:**
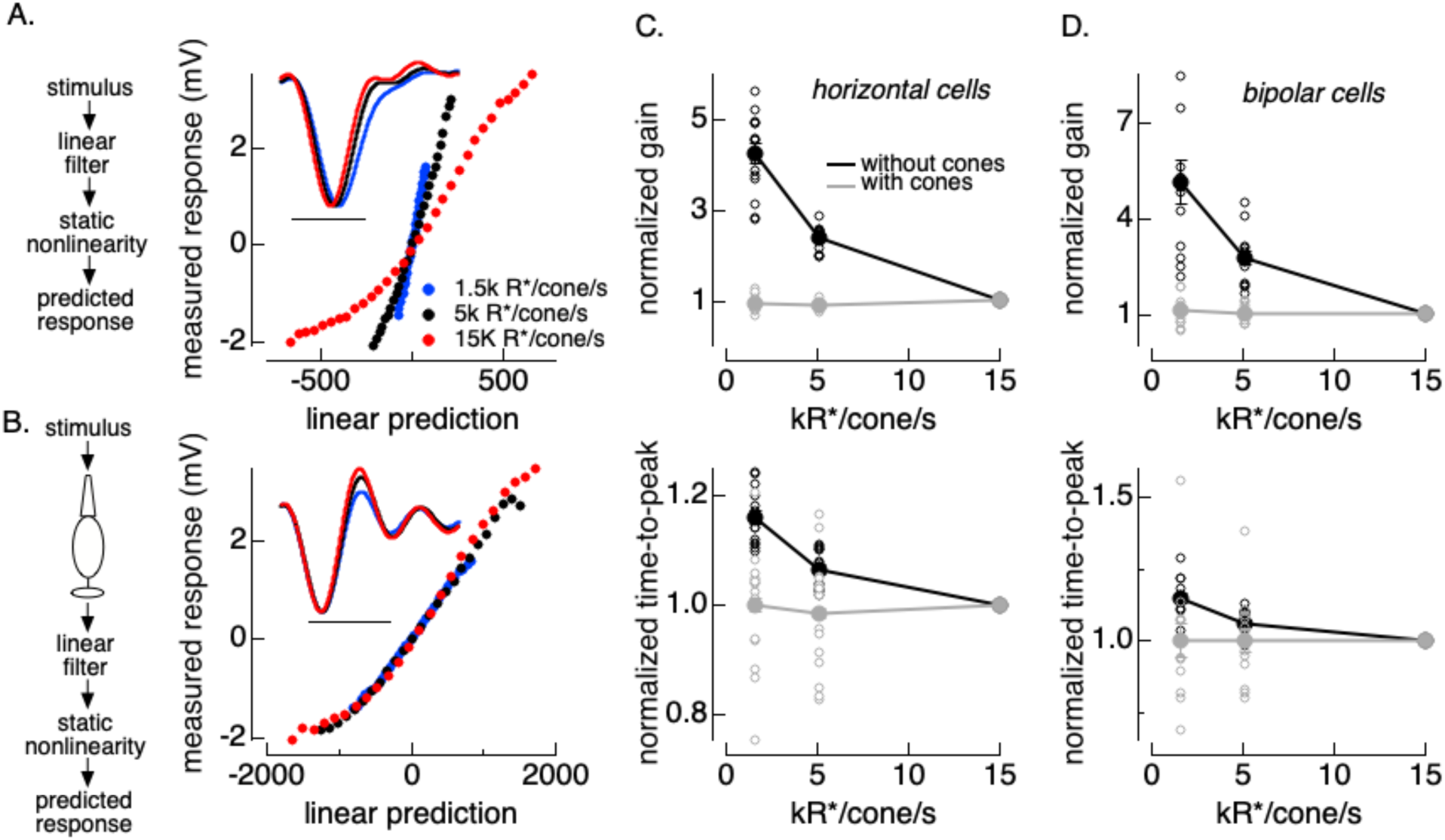
Effect of mean light level on linear-nonlinear models. A. Example horizontal cell. Standard linear-nonlinear model components at three mean light levels. B. As in A but for linear-nonlinear models that include an adapting cone front end. The output of the cone model provided the linear-nonlinear model input. C. Summary of gain (slope of nonlinearity, top) and filter time-to-peak (bottom) changes for horizontal cells without (black) and with (grey) cone front end (n=16). Gain and time-to-peak are normalized to values at 15k R*/cone/s. Open circles plot results from individual cells and solid circles plot mean ± SEM. D as in C for bipolar cell excitatory inputs (n=13).

The changes in gain and kinetics in Figure 2C and D are consistent with known adaptive properties of cone phototransduction (Angueyra and Rieke, 2013; Chen et al., 2024; Schneeweis and Schnapf, 1999). We tested the ability of cone adaptation to account for adaptation in horizontal and bipolar responses by repeating the linear-nonlinear analysis using models that incorporated a phototransduction “front end” consisting of a computational model fit to cone responses (Chen et al., 2024). This model was fixed, and did not introduce any new free parameters. Instead, the cone phototransduction model transforms light inputs to an estimate of the resulting cone photocurrents, including nonlinear properties of that transformation.

Incorporating cone phototransduction into linear-nonlinear models removed much of the dependence of both the linear filter kinetics and the nonlinearity slope on light intensity - i.e. filters and nonlinearities measured at different mean light levels were largely overlapping (Figure 2B). Across cells, the systematic changes in gain and kinetics were minimal and not statistically significant for models that incorporated the cone front end (Figure 2C, D, gray).

Figures 1 and 2 together identify two nonlinearities that contribute to horizontal and bipolar cell responses. Time-dependent nonlinearities in phototransduction account for most of the changes in gain and kinetics associated with changes in mean light level. An additional nonlinear mechanism at the cone output synapse also shapes responses to time-varying inputs at a constant mean light level. Horizontal cells showed little or no adaptation to changes in temporal contrast (Figure S1; (Rieke, 2001)).

### Models incorporating outer retina nonlinearities capture dynamic stimuli

The experiments above analyze responses in steady state. To test how well models incorporating these two types of nonlinearity work for more complex temporal stimuli, we measured responses to Gaussian noise stimuli that changed in mean intensity and/or contrast every 500-1000 ms. This stimulus emulates the rapid changes in mean and contrast that can occur following gaze shifts (Frazor and Geisler, 2006). Mean intensities and contrasts were chosen randomly in each time epoch, with values that spanned a 10-fold range of intensities and a 4-fold range of temporal contrasts. Figure 3 shows an example of the response of a horizontal cell to a short section of such a stimulus.

**Figure 3.**
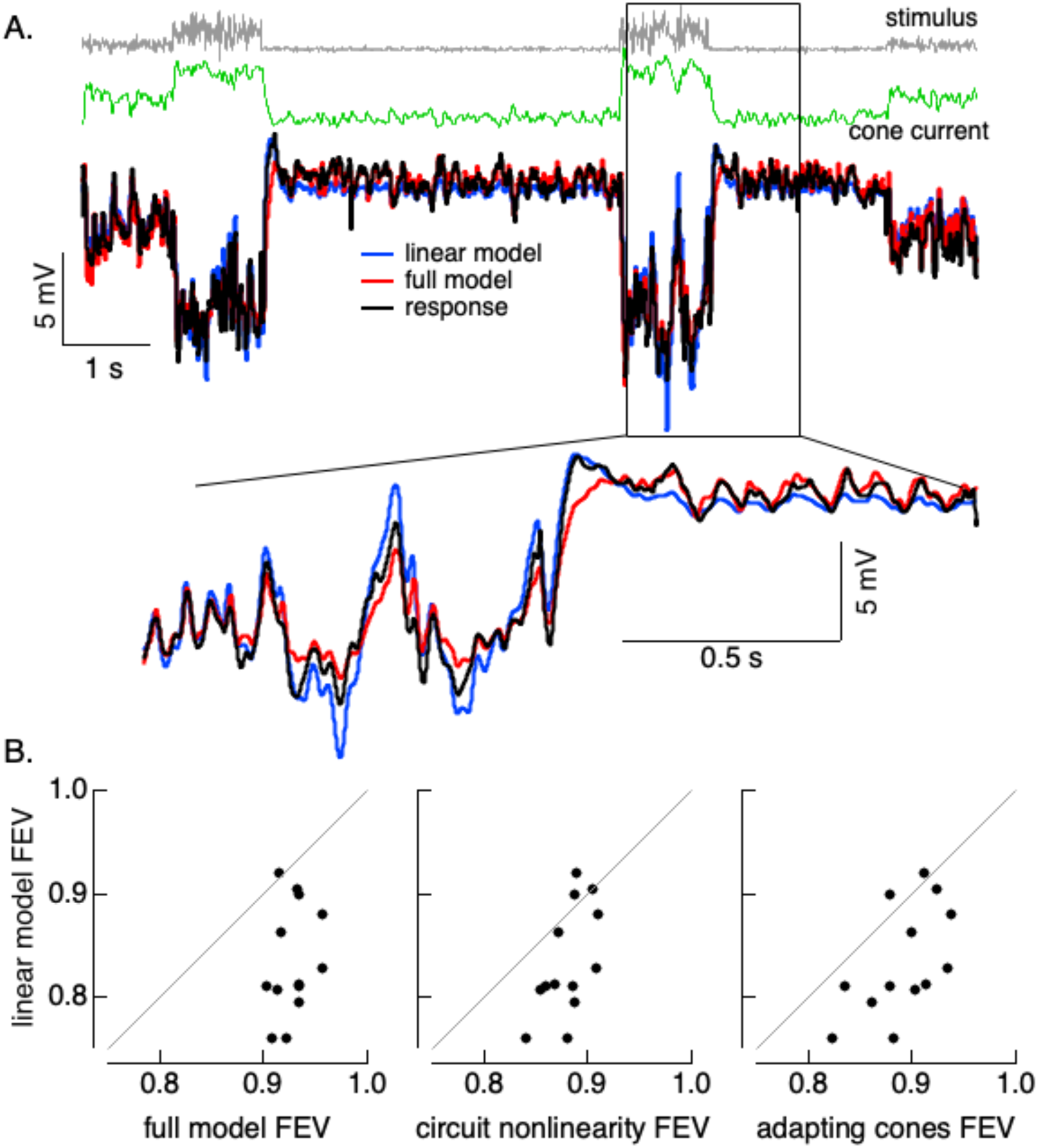
Responses and fits to variable mean noise stimulus. A. Short section of gaussian noise stimulus in which mean and contrast changes randomly every second (top), predicted cone photocurrents in response to this stimulus (second from top), and measured horizontal cell voltage and model predictions (bottom). B. Fraction explained variance for four different models. Differences in each panel were significant (left: p = 1e-5, middle: p = 8e-4, right: p = 1e-4, paired t-test). Mean light levels were 1500, 5000 and 15000 R*/cone/sec.

We fit these responses with four models: (1) a model incorporating a cone phototransduction front end followed by a nonlinear mechanism with different slopes for increments and decrements to account for the post-cone circuit nonlinearity (the “full model”); (2) a model in which the cones were linear but the circuit nonlinearity was retained (the “circuit nonlinearity” model); (3) a model with the cone front end without the circuit nonlinearity (the “adapting cones” model); and, (4) a fully linear model. We fit responses numerically (9 free parameters for models 1 and 2, 6 free parameters for models 3 and 4). The explained variance of the full model was > 90% for each of 13 horizontal cells, and this model performed substantially better than the fully linear model (Figure 3B, left). Performance of models lacking either the phototransduction or circuit nonlinearity fell between that of the full model and the linear model (Figure 3B, middle and right). The failures of the fully linear model can be seen in Figure 3A: the model overestimates responses at high light levels and underestimates them at low light levels.

Hence, for these dynamic stimuli both phototransduction and circuit nonlinearities contribute and the combined effect is a substantial improvement in the ability to predict measured responses.

### Horizontal cells integrate spatial inputs nonlinearly

How do the nonlinearities described above impact responses to spatial stimuli? The light responses of many retinal ganglion cells depend not only on the mean intensity of the light within their receptive field center, but also the distribution of light across space (Enroth-Cugell and Robson, 1966; Hochstein and Shapley, 1976). This dependence is due to nonlinear receptive field subunits, which cause light and dark regions of a stimulus not to cancel.

Nonlinear subunits are classically identified from responses to gratings in which the light and dark bars are balanced and the average intensity across space is equal to the background. A cell that sums stimuli across space linearly will not respond to such gratings, while cells with nonlinear subunits will respond since responses to the light and dark bars of the grating will not cancel (Figure 4A). Any nonlinearity that creates asymmetric responses to contrast increments and decrements (i.e. light and dark bars of the grating) suffices to generate responses to gratings; for a temporally-modulated grating, these responses occur at twice the modulation frequency and hence are often referred to as “F2” responses.

**Figure 4:**
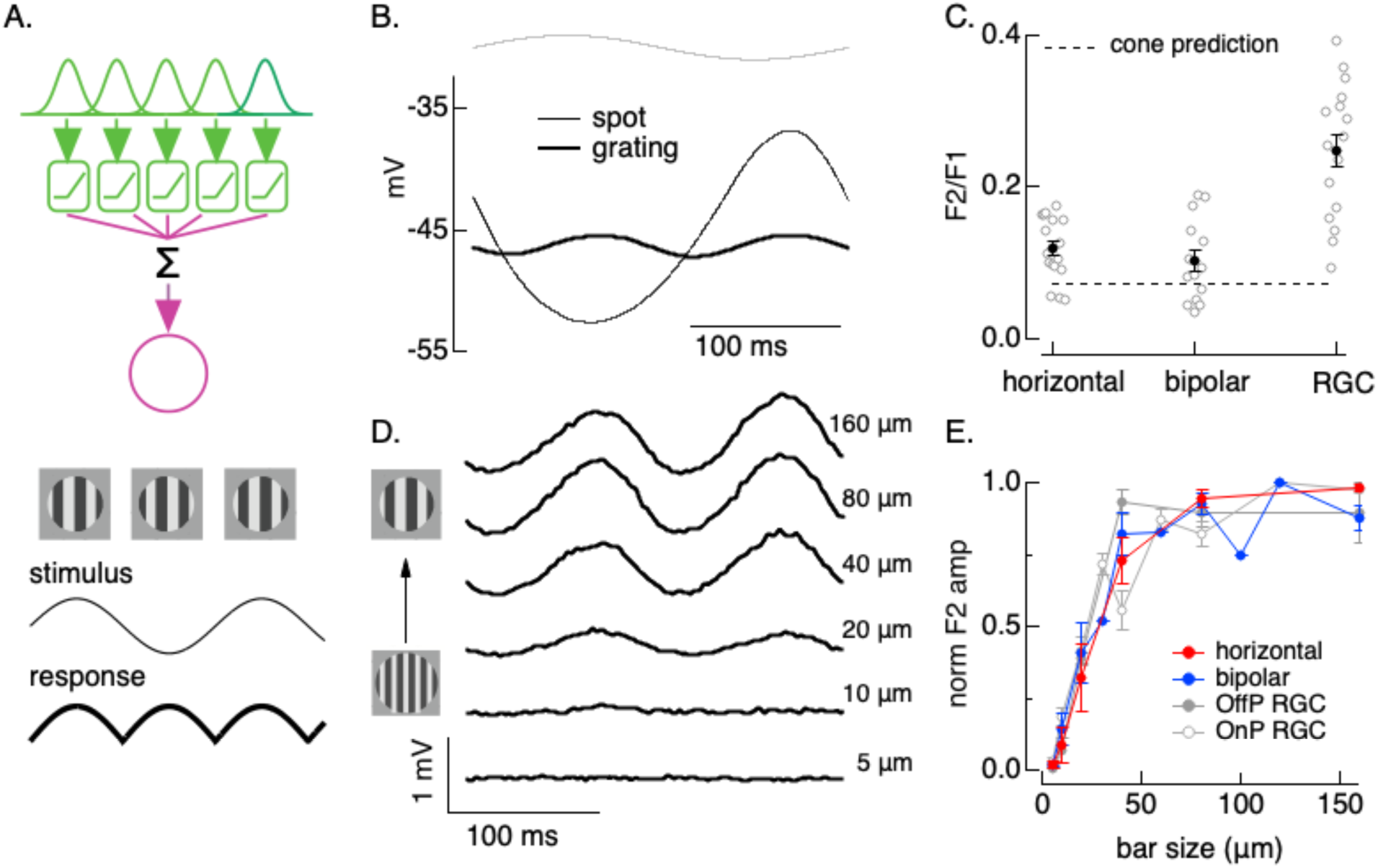
Nonlinear spatial integration in horizontal and bipolar cells. A. Logic behind the classic test for nonlinear spatial integration. Spatially local nonlinearities (green boxes) cause responses to light and dark bars of the grating to fail to cancel, resulting in a frequency doubled “F2” response (bottom) when the grating contrast is modulated sinusoidally (thin trace, second from bottom). B. Horizontal voltage responses to a modulated spot and grating. C. Strength of nonlinear response (quantified as amplitude of responses to the grating divided by the response to the spot) for horizontal cells (n=17), bipolar cells (n=14) and parasol RGCs (n=16). D. Dependence of horizontal response on grating bar width. Traces are displaced vertically. E. Summary (mean ± SEM) of dependence of nonlinear response on bar width for horizontal cells (n=4), bipolar cells (n=6) and parasol ganglion cells (n=11). All recordings at a mean light intensity of 15000 R*/cone/sec and all gratings had a contrast of 0.9.

Subunits have classically been attributed to nonlinear properties of bipolar synapses (Demb et al., 2001a), with the implicit assumption that cells in the outer retina integrate inputs across space linearly. Although cone photoreceptors respond to low contrast or rapidly varying inputs near-linearly (e.g. Figure 1), they respond nonlinearly to many high contrast stimuli due to rapid adaptation, resulting in larger responses to decrements than equal contrast increments (Angueyra et al., 2022; Chen et al., 2024). This suggests that cones could provide nonlinear subunits in the receptive fields of horizontal and bipolar cells. Consistent with this hypothesis, horizontal cells exhibited clear frequency-doubled responses to contrast-reversing gratings (Figure 4B). Excitatory synaptic inputs to bipolar cells exhibited similar frequency-doubled responses (not shown).

We quantified the strength of the nonlinear spatial responses from the ratio of the amplitude of the F2 response to that of the response to a spatially-uniform spot with the same contrast and temporal frequency as the grating (the “F1” response; Figure 4C). The strength of nonlinear spatial responses of horizontal and bipolar cells was about half that of responses in retinal ganglion cells (RGCs) measured under identical conditions (Figure 4C). Horizontal and bipolar nonlinear spatial integration was somewhat larger than predicted from increment/decrement asymmetries in cone phototransduction (dashed line in Figure 4C); this is consistent with a primary role of cone phototransduction and a secondary role of synaptic nonlinearities in generating the frequency-doubled responses.

The dependence of the frequency-doubled response on grating bar size provides an estimate of the size of the underlying receptive field subunits. Horizontal and bipolar F2 responses increased with bar size (Figure 4D), with a dependence similar to that measured in responses of parasol RGCs (Figure 4E). Horizontal F2 responses were apparent for gratings with bars as small as 10-20 μm - similar to the spacing between cone photoreceptors in the peripheral retina where we made our recordings.

Figure 4 shows that nonlinearities in the outer retina impact how horizontal and bipolar cells integrate spatial inputs. The spatial scale and strength of these responses suggests an origin in the cone photoreceptors. Further, these nonlinear spatial responses are similar in spatial scale and about half as strong as nonlinear spatial responses measured in RGCs; this suggests that the outer retina nonlinearities contribute to RGC responses, a topic we return to below.

### Nonlinear subunits shape horizontal cell responses to natural images

We next tested whether the subunit behavior identified with contrast reversing gratings shapes horizontal cell responses to natural inputs. To do this, we recorded responses to flashed natural images and corresponding spatially homogeneous “linear-equivalent discs” which provided an estimate of the expected response for a cell without nonlinear subunits (i.e. a cell that linearly summed inputs across space; (Turner and Rieke, 2016)). At the start of each recording, we estimated the receptive field by fitting the dependence of the response on spot size (Figure 5A). For each natural image, we then determined the contrast of a homogeneous disc – the linear-equivalent disc - that produced the same output as the image when passed through the estimated horizontal cell receptive field (see Methods). Linear spatial integration predicts that the linear-equivalent disc and image should produce identical responses; systematic differences in responses indicate nonlinear spatial integration.

**Figure 5:**
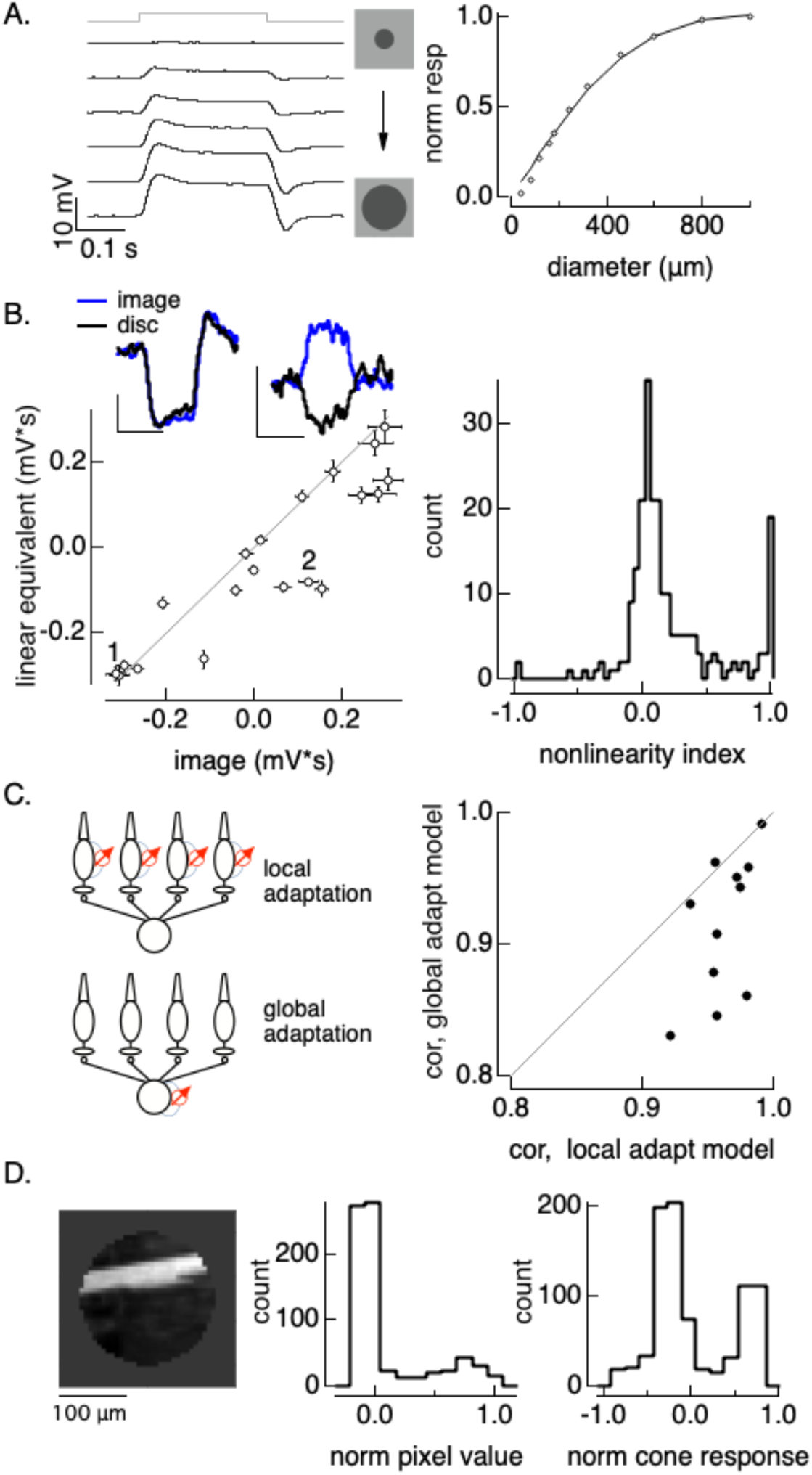
Nonlinear spatial integration impacts horizontal cell responses to natural images. A. (left) Receptive field measured by responses to flashed spots with increasing size from top to bottom. Traces are displaced vertically. (right) Integrated response as a function of spot diameter; solid line is a gaussian fit. B. (left) Responses to images and corresponding linear-equivalent discs. Insets at top show two highlighted examples, indicated by numbers in the main panel. (right) Nonlinearity index collected across images and cells. Responses to images different significantly from those to linear equivalent discs (p < 1e-10, paired t-test). C. Comparison of ability of local vs global adaptation to account for horizontal responses to flashed natural images. Correlation values for the local adaptation model were significantly higher than those for the global adaptation model (p = 0.006, paired t-test). D. Image 2 from panel B and histograms of stimulus pixel values (middle) and estimated cone responses (right). The mean pixel value is positive and the mean estimated cone response is negative for this image.

Responses to images differed systematically from those to the corresponding linear equivalent stimuli (Figure 5B). Specifically, responses to images were systematically less negative than those to linear equivalent discs (i.e. the points in Figure 5B lie on or to the right of the line of equality). These differences can be substantial - in some cases responses to the image and disc had opposite polarities (e.g. patch 2 in Figure 5B). We quantified this difference across images and cells using a nonlinearity index (NLI), which normalizes the difference in responses to the image and linear equivalent disc by the sum of the absolute values of the responses. An NLI of zero means that responses to the image and linear equivalent disc were equal, a positive NLI means that the response to the image was more positive than that to the disc (i.e. to the right of the line in Figure 5B), and a negative NLI means that the disc response was more positive than the image response (i.e. to the left of the line in Figure 5B). Measured NLIs were skewed to positive values (Figure 5B, right), consistent with the trend from the example cell in Figure 5B. Further, many images had NLIs near 1 (image 2 in Figure 5C is one example), indicating that the response differences were often as large as the responses themselves. This analysis indicates that spatial structure in natural images impacts horizontal cell responses and that this impact can be substantial.

Past work shows that cone responses to contrast increments and decrements are asymmetrical due the same adaptive mechanisms invoked to explain horizontal cell responses in Figures 2-4 (Angueyra et al., 2022; Chen et al., 2024). Can this asymmetry explain the sensitivity of horizontal cell responses to spatial structure? To answer this question, we predicted responses to natural image patches using the cone phototransduction model, which accounts for the increment/decrement asymmetry (see Methods). We linearly summed predicted cone responses across space using the horizontal cell’s estimated receptive field. We compared this model - in which adaptation occurs locally in each cone - with a model in which adaptation occurs after combining signals across cones (the “global nonlinearity” model). This comparison tests specifically whether adaptation needs to operate locally to account for the horizontal sensitivity to spatial structure. The local adaptation model predicted measured responses with higher accuracy than the global adaptation model, with correlations between predicted and measured responses exceeding 0.9 across 11 cells and images (Figure 5C, right). Adding a post-transduction synaptic nonlinearity did not substantially improve predictions, consistent with a dominant role of transduction in horizontal cell nonlinear spatial integration (see also Figure 4C).

How can the incorporation of cone adaptation explain the opposite polarity responses such as image 2 in Figure 5B? Figure 5D (right) shows that specific image, masked to the region presented during the experiment to fill the horizontal receptive field center. The majority of the pixels in the image have small negative contrasts (Figure 5D, middle), with a subset of pixels with large positive contrasts; these are the pixels comprising the bright branch. The average pixel value is positive, and correspondingly the linear equivalent disc contrast is positive and it hyperpolarizes the cell (Figure 5B, top right inset, black trace). Rapid adaptation in the cone phototransduction cascade causes the distribution of predicted cone photocurrents to differ from the pixel distribution - specifically, dark pixels are associated with a higher phototransduction gain than bright pixels (Figure 5D, right). This in turn causes the mean photocurrent to be negative, providing an explanation for why the image depolarizes the cell (Figure 5B, top right inset, blue trace). Similar shifts in the pixel distribution underlie the other large positive nonlinearity indices in Figure 5B.

The experiments and analysis summarized in Figure 5 show that nonlinear spatial integration contributes substantially to horizontal cell responses to natural images, and that these contributions can be largely accounted for by local adaptation in cone phototransduction.

### Nonlinearities shape horizontal cell responses to natural movies

The results above identify clear nonlinearities that contribute to horizontal cell responses to temporal and spatial stimuli. Here we evaluate the impact of these nonlinearities on responses to natural movies.

We first experimentally tested the importance of nonlinear spatial integration in shaping responses to natural movies by generalizing the linear-equivalent disc stimuli used in the experiments of Figure 5 to movies (Turner and Rieke, 2016). We compared responses to two movies: (1) a natural movie consisting of a fixed image moved with a trajectory corresponding to measured human eye movements (Van Der Linde et al., 2009); (2) a movie in which each video frame of the original movie was replaced with its corresponding linear-equivalent disc (computed as in Figure 5; see Methods). A cell that integrates linearly across space should respond near-identically to the two movies. Responses to the two movies instead differed substantially (Figure S1). These differences motivated a more complete analysis of responses to natural movies.

We compared predictions of responses to natural movies using four different models (Figure 6A; see Methods for details). These are the same models used in Figure 3 with the addition of a spatial component. Each started with a model for the conversion of light inputs to cone photocurrents. That consisted of either a full (nonlinear) model for cone phototransduction or a matched linear transduction model. The transduction models were fixed and did not introduce any free parameters. The transduction stage was followed by a synaptic stage, which was either linear or differentially weighted positive and negative changes in cone photocurrents. The nonlinear synaptic stage introduced one free parameter (the difference in gain applied to increments vs decrements). Cone outputs were then summed by a two-dimensional Gaussian receptive field. Models with the local synaptic nonlinearity had 9 free parameters, models without it had 8 free parameters.

**Figure 6:**
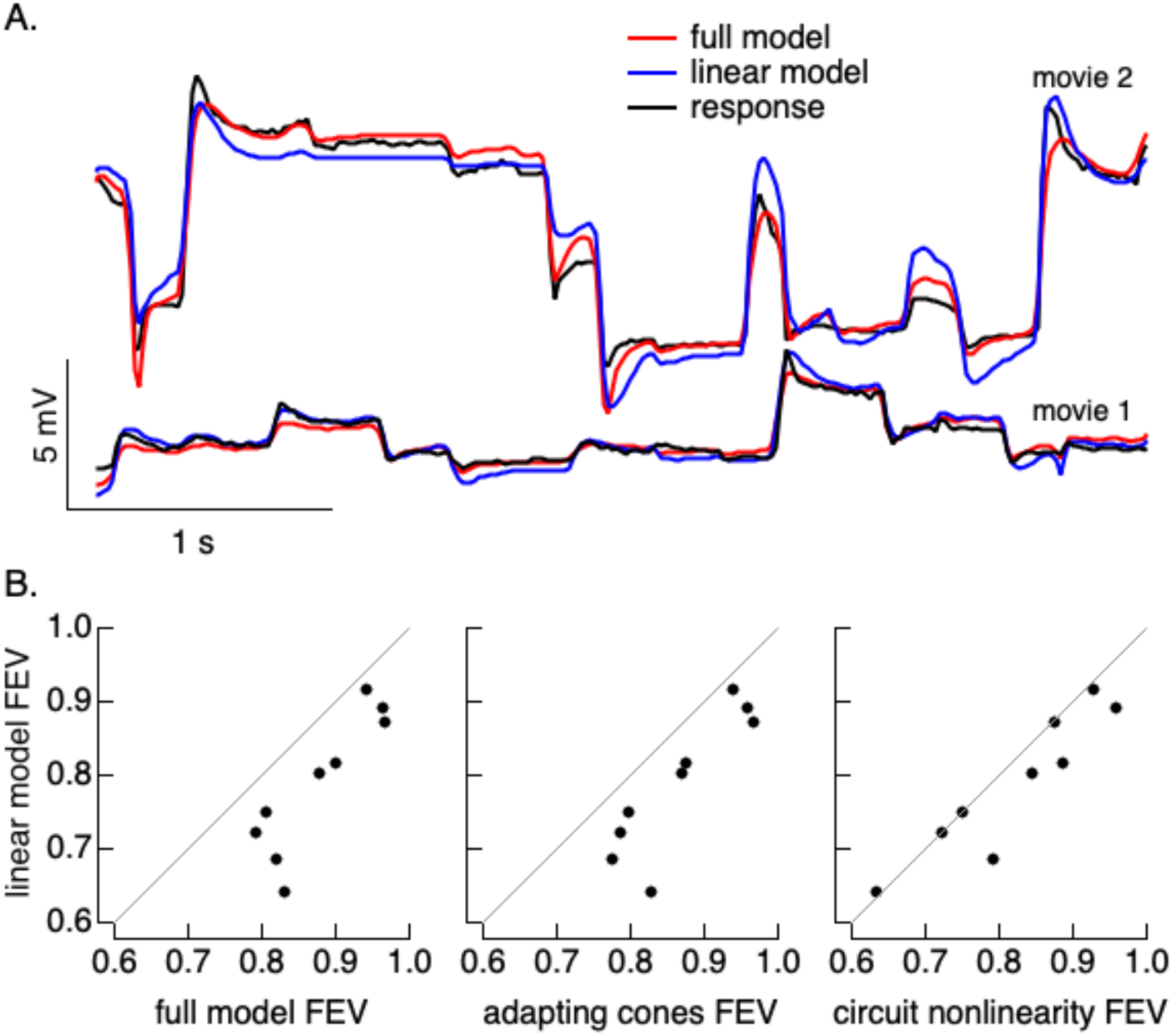
Nonlinear model fits to horizontal cell responses to natural movies. A. Responses to two example natural movies and fits of linear and nonlinear models. B. Fraction explained variance for four models with different combinations of cone phototransduction and a post-cone circuit nonlinearities. Differences in each panel were significant (left: p=5e-4, middle: p = 1e-3, right: p = 0.04, paired t-test).

Models that included both phototransduction and post-transduction local nonlinearities made more accurate predictions than linear models (Figure 6B, left). Models with one of the two nonlinear mechanisms alone also outperformed linear models (Figure 6B, middle and right). The majority of the performance gain from the full nonlinear model was also captured by a model with nonlinear cone phototransduction alone. This analysis indicates that outer retinal nonlinearities, especially nonlinearities in cone phototransduction, shape horizontal cell responses to natural movies.

### Outer retina nonlinearities cause receptive fields to dynamically shift with changes in context

What impact do the outer retina nonlinearities described above have on retinal output signals? While we cannot answer this question in general here, the following sections provide two examples of how the mechanisms described above impact ganglion cell receptive fields.

We start by testing the impact of outer retina nonlinearities on context dependence of the receptive field (Goldin et al., 2022). RGC receptive fields differ when measured in the presence and absence of a context-defining image. Spatially local adaptation should contribute to this context dependence by increasing the weighting of stimuli in spatial regions with high gain and decreasing the weighting of regions with low gain (Figure 7A, top). In the example of Figure 7A, this should shift the effective receptive field to the left towards the dark region of the context-defining image.

**Figure 7:**
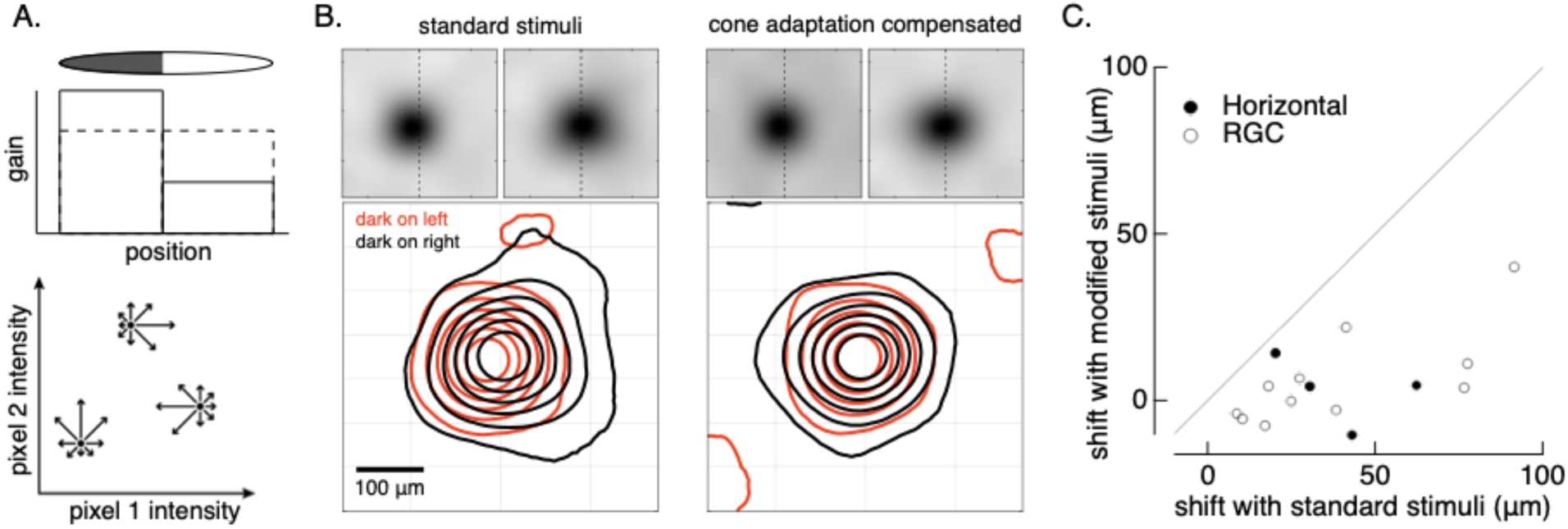
Context dependence of horizontal cell receptive fields. A. (top) Local adaptation (e.g. in the cone photoreceptors) is expected to shift the effective receptive field towards dark regions of the context defining stimulus. (bottom) The receptive field in the presence of a context-defining stimulus can be measured by adding multiple random perturbations to the stimulus, weighting each by the response it elicits and averaging. B. Horizontal cell receptive field measured with standard stimuli (left) and with stimuli in which the contrast has been adjusted in each pixel to compensate for adaptation in the cones (right). C. Summary of receptive field shifts across RGCs and horizontal cells. Shifts were estimated by computing the center-of-mass of the receptive field for each of the two grating conditions. p = 0.003 for RGCs (paired t-test).

To test this prediction, we added random spatial noise to a disc with a dark and bright half. We estimated the receptive field in presence of the disc by weighting several hundred noise stimuli by the response each elicited and averaging these weighted noise images (Figure 7A, bottom). This process measures the direction in which a cell’s response changes most rapidly given starting point given by the context-defining image. We repeated this process with the dark region of the disc either on the left or right. As predicted from local adaptation, the horizontal cell receptive field shifted towards the dark region of the disc (Figure 7B, left). To test whether this shift was consistent with cone adaptation, we adjusted the standard deviation of the spatial noise in the dark and bright regions to compensate for the expected effects of cone adaptation. This meant decreasing the magnitude of the noise in the dark region where the gain of cone responses is higher and increasing it in the bright region where the gain of cone responses is lower. The predicted gain changes - and corresponding changes in noise to compensate the gain changes - came directly from our cone model. Receptive fields computed for stimuli that compensated for the predicted cone adaptation lost most of their context dependence (Figure 7B, right).

Ganglion cell receptive fields exhibited a similar context dependence. We summarized these receptive field shifts by measuring the distance between the center-of-mass of the receptive fields measured in the presence of the two gratings. Receptive fields of both horizontal cells and ganglion cells shifted significantly for standard stimuli (x-axis in Figure 7C), and these changes decreased substantially for stimuli that compensated for expected cone adaptation (y-axis in Figure 7C).

In sum, these experiments show that outer retina nonlinearities contribute substantially to an unexpected dependence of RGC responses on the spatial context in which they are measured.

### Outer retina nonlinearities account for nonlinear RGC surrounds

We next explored the impact of outer retina nonlinearities on retinal ganglion cell receptive field surrounds. Activity of the receptive field surround depends on both the mean luminance and spatial contrast (Cook and McReynolds, 1998; Demb et al., 1999). Further, surrounds in the primate retina originate at least in large part from horizontal cells (Davenport et al., 2008; McMahon et al., 2004). This leads to the hypothesis that the nonlinear spatial properties of horizontal cells, as detailed in Figure 4, account for the sensitivity of the surround to spatial structure. Our recent work supports this hypothesis (Hong and Rieke, unpublished). We provide an additional test here.

We measured responses to asymmetric gratings that had different light and dark bar contrasts. Spatial structure depolarizes horizontal cells (Figures 4 and 5), while positive mean luminance hyperpolarizes them (Hong and Rieke, unpublished). This suggests that there is a balance point in which a spatially structured stimulus such as a grating with a net positive mean luminance will create little or no horizontal cell response because the impact of the stimulus mean and spatial structure will cancel; if horizontal cells dominate RGC surrounds, this same balance point should minimize the RGC surround response. This was indeed the case (Figure 7): gratings in which the dark bars had a contrast that was ∼40% of the bright bar contrast produced little or no change in horizontal cell voltage (Figure 7A) and the same gratings confined to On parasol RGC surrounds elicited little of no response (Figure 7B). The similarity of horizontal cell and On parasol surround responses held across the full range of gratings probed (Figure 7C). Further, the increment/decrement asymmetry in cone phototransduction accurately predicted both horizontal and RGC responses to the full collection of asymmetric grating stimuli (solid line in Figure 7C).

**Figure 8:**
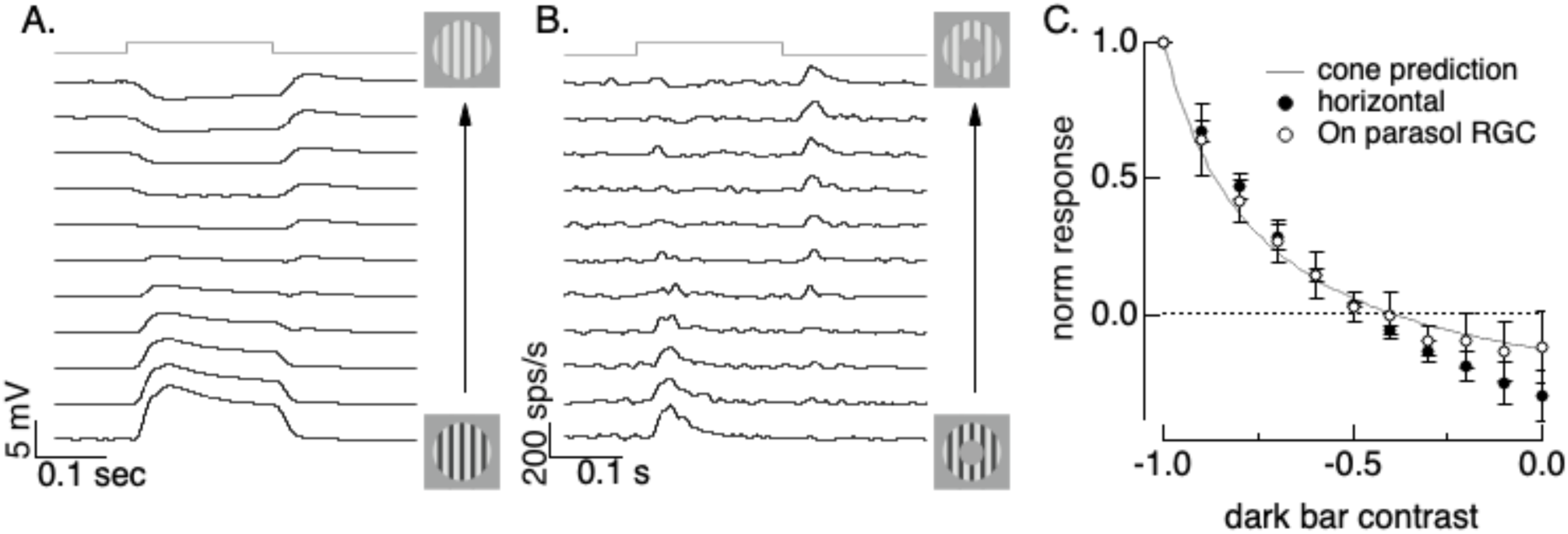
A. Horizontal responses to unbalanced gratings. The bottom trace shows the response to a standard flashed grating (equal light and dark bars) - then as you go up the dark bar contrast decreases to 0.9, 0.8, … to 0 in the top trace. B. As in A for On parasol spike response. Firing rate is an average of 6 trials of each stimulus. Stimuli are the same as A except for the mask covering the receptive field center. C. Measured and predicted responses (mean ± SD) to onset of gratings from On parasol (n=4) and horizontal cells (n=3). Responses are integrated across the time the grating is present. Predictions are from the cone model.

These results are consistent with a dominant role of horizontal cells in generating On parasol receptive field surrounds, and a dominant role of cone adaptation in generating surround sensitivity to spatial structure.

## Discussion

### Summary

We have characterized responses of the outer retina to a range of stimuli to determine how visual inputs are transformed through the first steps of the visual process. Two nonlinear mechanisms together provided an accurate description of responses across the range of stimuli probed: (1) adaptive nonlinearities in cone phototransduction, and (2) an additional nonlinear step at the cone output synapse. These nonlinearities shaped both horizontal cell and RGC responses to naturalistic stimuli.

### Limitations

We focused on responses of horizontal cells since they provide a means to characterize outer retinal signals without confounding impacts from inner retinal mechanisms. We collected a limited set of data from bipolar cells to verify that key properties of the horizontal cell responses were shared by bipolar cells. We did not compare different bipolar cell types as unambiguously identifying cell types in the flat mount recordings that we used primarily here is difficult.

We found that outer retinal mechanisms contribute to subunits of horizontal and RGC RFs and the signaling properties created by those subunits. This does not preclude inner retinal mechanisms such as nonlinearities at the bipolar output synapse that also contribute to subunits in the RGC RFs; indeed, subunit strength in the RGCs was stronger than that in horizontal and bipolar cells (Figure 4). Evaluating the relative contributions of these two sources of subunits to retinal outputs is difficult because they have similar properties.

We chose models with a limited number of free parameters so that model components could be interpreted directly in terms of circuit mechanisms. These models captured 80-90% of the variance of the horizontal responses to natural movies but also systematically missed some aspects of these responses. With longer recordings these systematic errors might guide improvements in model architecture, but we did not have enough data here to test this possibility.

### Origin of nonlinear receptive field subunits

The origin and functional impact of nonlinear RF subunits has been an important theme in work in the retina and early visual areas for many years. Subunits and the resulting nonlinear spatial integration have long been used to differentiate different RGC types, e.g. X and Y cells in cat (Enroth-Cugell and Robson, 1966) and midget and parasol cells in primate (Benardete and Kaplan, 1999a, 1999b; Kaplan and Shapley, 1986). Nonlinear subunits also play an important role in many specialized retinal computations such as sensitivity to various forms of motion and spatial texture (Baccus et al., 2008; Kuo et al., 2016; Manookin et al., 2018).

Subunits are created by nonlinear mechanisms that operate on a spatial scale finer than the RGC RF. These local nonlinearities cause RGC responses to depend on spatial structure rather than just the integrated light intensity within their RF. Such sensitivity is present in RGC excitatory synaptic inputs, and this observation suggests that the bipolar cell output synapse provides the nonlinearity creating subunits (Demb et al., 2001a, 1999). The origin is important computationally since the order of linear and nonlinear operations matters - e.g. a circuit in which excitatory and inhibitory inputs converge followed by a shared nonlinearity is not equivalent to one with separate nonlinearities for each input type followed by convergence.

Here we show that responses of horizontal and bipolar cells show clear subunit properties due to nonlinearities generated within the cone photoreceptors. Horizontal cell responses to a variety of stimuli were quantitatively accounted for by models that incorporated nonlinearities in cone phototransduction and the cone output synapse. The similarity of the subunits measured in horizontal cells with those in RGCs suggests that they contribute to nonlinear spatial integration by RGCs. Subunits originating in the outer retina should be shared by all downstream retinal neurons. Hence, together with previous work in salamander (Schreyer and Gollisch, 2021), our findings indicate that subunits originate at least in part earlier than generally appreciated, prior to many of the mechanisms that differentiate signals in different parallel retinal pathways. This differs from the architecture of most empirical models for retinal signaling, which begin with a linear pooling over space and emphasize nonlinear mechanisms that differ across different parallel pathways. The distinction between early and late nonlinearities has an important impact on models for retinal computations (Gollisch and Meister, 2010).

### Functional roles of horizontal cells

Horizontal cells shape retinal responses in several important ways. In space, lateral inhibition from horizontal cells contributes to center-surround receptive fields that enhance contrast edges (Barlow, 1953; Kuffler, 1953; Marr and Hildreth, 1980). This mechanism appears to dominate RF surrounds in the primate retina (Davenport et al., 2008; McMahon et al., 2004). In time, horizontal feedback to cones can sharpen temporal responses by suppressing low temporal frequency components of the cone response (Baylor et al., 1971) (see also (Chapot et al., 2017). Linear or quasi-linear models are often used to account for these aspects of horizontal cell responses (Verweij et al., 1999).

Receptive field surrounds are also engaged by spatially-structured inputs (Cook and McReynolds, 1998; Demb et al., 1999). We found recently that the sensitivity of the RF surround to spatial structure in the primate retina originates largely from horizontal cells (Hong and Rieke, unpublished). This causes Off RGCs to respond preferentially to contrast edges such as those created by discontinuities in texture, while On RGCs respond preferentially to spatially-extended textures. Consistent with that work, we show here that horizontal cell responses to spatial structure share several properties with nonlinear responses of RGC surrounds.

The spatially local adaptive nonlinearities that create horizontal cell subunits also contribute to context dependence of the RGC RFs (Goldin et al., 2022). One important piece of evidence for this conclusion came from stimuli designed to negate outer retina nonlinearities. In the future, extending the work here to provide a complete quantitative model for horizontal cell responses could permit other causal manipulations to probe their role in shaping retinal output and responses of downstream visual neurons.

## Methods

Primate retinas (*Macaca nemestrina, Macaca mulatta and Macaca fascicularis*) were obtained through the Tissue Distribution Program of the National Primate Research Center at the University of Washington. Procedures adhered to ethical guidelines approved by the University of Washington Institutional Animal Care and Use Committee (IACUC). We did not observe any substantial differences from retinas were obtained from animals of either sex.

Retinas were prepared and stored following published procedures (Turner and Rieke, 2016). For most experiments, small pieces of retina (∼2×3 mm) with the pigment epithelium attached were mounted RGC side up on poly-L-lysine-coated coverslips. Experiments were performed on peripheral retina (20–50° from the fovea), with eccentricity determined based on cone density.

The same preparation was used for recordings from horizontal cells, bipolar cells and RGCs. Some horizontal and bipolar recordings were made in retinal slices, prepared as previously described (Grimes et al., 2014).

### Electrophysiological Recordings

Retinas were superfused continuously with oxygenated Ames’ medium (pH ∼7.4, bubbled with 95% O₂/5% CO₂) at 33-34 °C. Data were acquired at 10 kHz using a Multiclamp 700B amplifier (Molecular Devices), filtered at 3 kHz (900-CT, Frequency Devices), digitized with an ITC-18 analog-to-digital board (HEKA Instruments), and recorded using the Symphony data acquisition software package (http://symphony-das.github.io).

The health and sensitivity of each flat-mount preparation was tested by delivering a uniform, 5% contrast, 4 Hz modulated stimulus. Data was collected only from retinas in which this stimulus produced at least a 20 sps/s modulation of spike rate in On parasol RGCs. This same quality control applied regardless of the cell types form which data was collected.

Extracellular recordings from RGCs were performed using borosilicate glass pipettes filled with Ames’ medium. Current-clamp recordings from horizontal cells used intracellular solutions containing 123 mM K-aspartate, 10 mM KCl, 10 mM HEPES, 1 mM MgCl₂, 1 mM CaCl₂, 2 mM EGTA, 4 mM Mg-ATP, and 0.5 mM Tris-GTP, pH 7.3. Voltage-clamp recordings from bipolar or horizontal cells used intracellular solutions containing 105 mM Cs-methanesulfonate, 10 mM TEA-Cl, 20 mM HEPES, 10 mM EGTA, 2 mM QX-314, 5 mM Mg-ATP, and 0.5 mM Tris-GTP, pH ∼7.3 with CsOH. Pipette resistances were 8-12 MΩ for bipolar and horizontal cell recordings. Access resistance was < 20 MΩ for bipolar recordings and horizontal cell recordings.

Horizontal cells were distinguished by their soma location at the border between the inner nuclear layer and outer plexiform layer, their large receptive fields and the characteristic kinetics of their light responses (e.g. Figure 5A). Bipolar cells were identified based on soma location in the inner nuclear layer, their small receptive fields, and sustained responses to 0.5 sec light steps.

### Visual Stimuli

Spatially uniform visual stimuli were delivered using a 405 nm LED that uniformly illuminated a 500 um diameter spot at the retina. Spatially-varying visual stimuli were generated using the Stage software package (http://stage-vss.github.io) and displayed on either a LightCrafter 4500 projector or an eMagin OLED display. In all cases light was focused on the photoreceptors through a 10x water-immersion microscope objective.

### Linear equivalent stimuli

The linear equivalent stimuli used to probe spatial integration used an estimate of a cell’s receptive field to weight each pixel of the stimulus and determine the contrast *C* of a spatially homogeneous disc predicted to elicit the same response as the original image

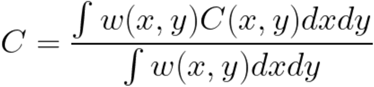

where *w(x,y)* is the RF weight and *C(x,y)* is the image pixel contrast at each spatial location. The RF was assumed to be Gaussian, fit to each cell’s measured response-vs-area curve as in Figure 5A. This process was applied to each video frame for the natural movies in Figure 6.

### Modeling

Several models used an initial step in which stimulus intensities were converted to estimated cone transduction currents. This conversation used the cone phototransduction model built into isetbio (http://isetbio.org/; see also (Chen et al., 2024)). The local adaptation model in Figure 5 used a simpler cone front-end that incorporated only a gain term, derived from a Weber curve describing the relationship between cone sensitivity and mean light intensity (half-desensitizing background of 2000 R*/cone/s; (Angueyra and Rieke, 2013; Cao et al., 2014)). Spatially-local nonlinearities in models used to fit natural movies (Figure 6) applied a different scaling factor to positive and negative contrast pixels. Models with a more complex local nonlinearity (a sigmoid) did not improve performance, and hence the simpler model with fewer free parameters was used.

## Acknowledgements

We thank S. Cunnington for technical support and the Tissue Distribution Program (NIH grant OD010425) at the Washington National Primate Research Center, particularly C. English and A. Baldessari, for providing tissue. We also thank Samuele Virgili for valuable feedback on the manuscript. This work was supported by NIH grants EY028542 and EY036594 and a Stein Foundation Innovation Award.

## Supplementary Figures

**Figure S1:**
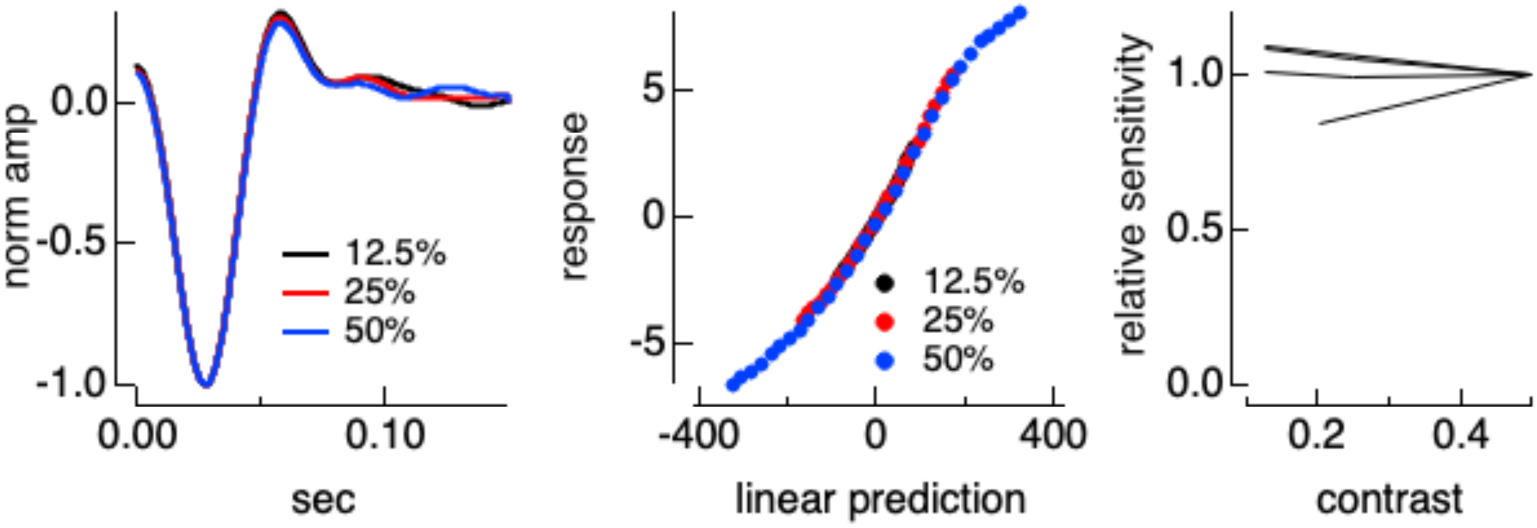
Little or no contrast adaptation. LN model fits to responses of a horizontal cell to Gaussian noise at the three contrasts indicated. Right panel shows the sensitivity, measured from the slope of the nonlinearity, for three such experiments.

**Figure S2:**
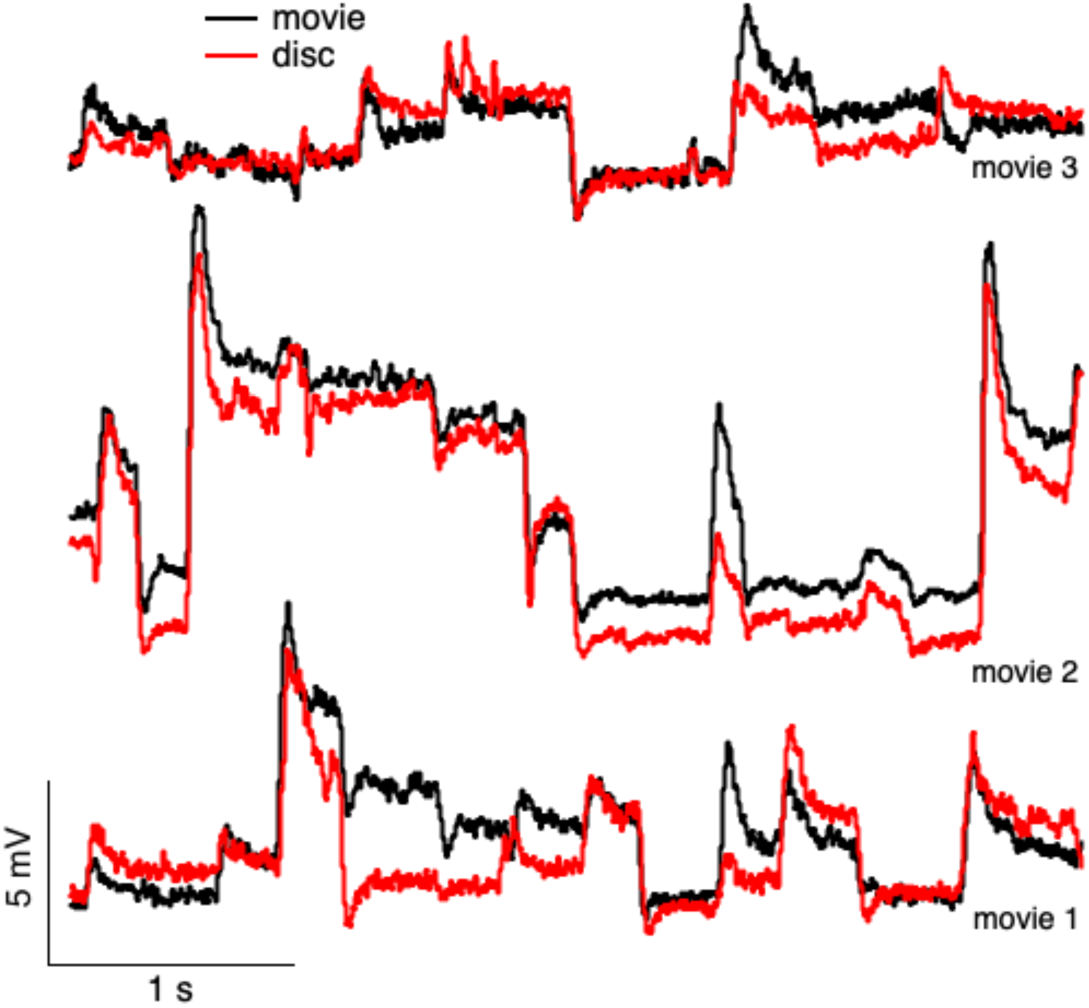
Horizontal cell responses to linear equivalent (red) and Doves (black) movies.

## References

Adelson, E.H., Bergen, J.R., 1985. Spatiotemporal energy models for the perception of motion. J. Opt. Soc. Am. A 2, 284–299. 10.1364/josaa.2.000284

Angueyra, J.M., Baudin, J., Schwartz, G.W., Rieke, F., 2022. Predicting and Manipulating Cone Responses to Naturalistic Inputs. J. Neurosci. Off. J. Soc. Neurosci. 42, 1254–1274. 10.1523/JNEUROSCI.0793-21.2021

Angueyra, J.M., Rieke, F., 2013. Origin and effect of phototransduction noise in primate cone photoreceptors. Nat Neurosci 16, 1692–1700.

Baccus, S.A., Olveczky, B.P., Manu, M., Meister, M., 2008. A retinal circuit that computes object motion. J Neurosci 28, 6807–6817.

Barlow, H.B., 1953. Summation and inhibition in the frog’s retina. J Physiol 119, 69–88.

Baylor, D.A., Fuortes, M.G., O’Bryan, P.M., 1971. Receptive fields of cones in the retina of the turtle. J. Physiol. 214, 265–294. 10.1113/jphysiol.1971.sp009432

Benardete, E.A., Kaplan, E., 1999a. The dynamics of primate M retinal ganglion cells. Vis Neurosci 16, 355–368.

Benardete, E.A., Kaplan, E., 1999b. Dynamics of primate P retinal ganglion cells: responses to chromatic and achromatic stimuli. J Physiol 519 Pt 3, 775–790.

Cao, L.H., Luo, D.G., Yau, K.W., 2014. Light responses of primate and other mammalian cones. Proc Natl Acad Sci U A 111, 2752–2757.

Chapot, C.A., Behrens, C., Rogerson, L.E., Baden, T., Pop, S., Berens, P., Euler, T., Schubert, T., 2017. Local Signals in Mouse Horizontal Cell Dendrites. Curr. Biol. 27, 3603–3615.e5. 10.1016/j.cub.2017.10.050

Chen, Q., Ingram, N.T., Baudin, J., Angueyra, J.M., Sinha, R., Rieke, F., 2024. Predictably manipulating photoreceptor light responses to reveal their role in downstream visual responses. eLife 13, RP93795. 10.7554/eLife.93795

Cook, P.B., McReynolds, J.S., 1998. Lateral inhibition in the inner retina is important for spatial tuning of ganglion cells. Nat Neurosci 1, 714–719.

Davenport, C.M., Detwiler, P.B., Dacey, D.M., 2008. Effects of pH buffering on horizontal and ganglion cell light responses in primate retina: evidence for the proton hypothesis of surround formation. J Neurosci 28, 456–464.

Demb, J.B., Haarsma, L., Freed, M.A., Sterling, P., 1999. Functional circuitry of the retinal ganglion cell’s nonlinear receptive field. J Neurosci 19, 9756–9767.

Demb, J.B., Zaghloul, K., Haarsma, L., Sterling, P., 2001a. Bipolar cells contribute to nonlinear spatial summation in the brisk-transient (Y) ganglion cell in mammalian retina. J Neurosci 21, 7447–7454.

Demb, J.B., Zaghloul, K., Sterling, P., 2001b. Cellular basis for the response to second-order motion cues in Y retinal ganglion cells. Neuron 32, 711–721.

Endeman, D., Kamermans, M., 2010. Cones perform a non-linear transformation on natural stimuli. J Physiol 588, 435–446.

Enroth-Cugell, C., Robson, J.G., 1966. The contrast sensitivity of retinal ganglion cells of the cat. J Physiol 187, 517–552.

Field, G.D., Rieke, F., 2002. Nonlinear signal transfer from mouse rods to bipolar cells and implications for visual sensitivity. Neuron 34, 773–785.

Frazor, R.A., Geisler, W.S., 2006. Local luminance and contrast in natural images. Vis. Res 46, 1585–1598.

Goldin, M.A., Lefebvre, B., Virgili, S., Pham Van Cang, M.K., Ecker, A., Mora, T., Ferrari, U., Marre, O., 2022. Context-dependent selectivity to natural images in the retina. Nat. Commun. 13, 5556. 10.1038/s41467-022-33242-8

Gollisch, T., Meister, M., 2010. Eye smarter than scientists believed: neural computations in circuits of the retina. Neuron 65, 150–164.

Gollisch, T., Meister, M., 2008. Rapid neural coding in the retina with relative spike latencies. Science 319, 1108–1111.

Grimes, W.N., Hoon, M., Briggman, K.L., Wong, R.O., Rieke, F., 2014. Cross-synaptic synchrony and transmission of signal and noise across the mouse retina. Elife 3.

Heeger, D.J., Simoncelli, E.P., Movshon, J.A., 1996. Computational models of cortical visual processing. Proc. Natl. Acad. Sci. U. S. A. 93, 623–627. 10.1073/pnas.93.2.623

Hochstein, S., Shapley, R.M., 1976. Quantitative analysis of retinal ganglion cell classifications. J Physiol 262, 237–264.

Kaplan, E., Shapley, R.M., 1986. The primate retina contains two types of ganglion cells, with high and low contrast sensitivity. Proc Natl Acad Sci U A 83, 2755–2757.

Kuffler, S.W., 1953. Discharge patterns and functional organization of mammalian retina. J Neurophysiol 16, 37–68.

Kuo, S.P., Schwartz, G.W., Rieke, F., 2016. Nonlinear spatiotemporal integration by electrical and chemical synapses in the retina. Neuron in press.

Liu, B., Hong, A., Rieke, F., Manookin, M.B., 2021. Predictive encoding of motion begins in the primate retina. Nat. Neurosci. 24, 1280–1291. 10.1038/s41593-021-00899-1

Manookin, M.B., Patterson, S.S., Linehan, C.M., 2018. Neural Mechanisms Mediating Motion Sensitivity in Parasol Ganglion Cells of the Primate Retina. Neuron 97, 1327–1340.e4. 10.1016/j.neuron.2018.02.006

Marr, D., Hildreth, E., 1980. Theory of edge detection. Proc. R. Soc. Lond. B Biol. Sci. 207, 187–217. 10.1098/rspb.1980.0020

McMahon, M.J., Packer, O.S., Dacey, D.M., 2004. The classical receptive field surround of primate parasol ganglion cells is mediated primarily by a non-GABAergic pathway. J Neurosci 24, 3736–3745.

Rieke, F., 2001. Temporal contrast adaptation in salamander bipolar cells. J Neurosci 21, 9445–9454.

Sampath, A.P., Rieke, F., 2004. Selective Transmission of Single Photon Responses by Saturation at the Rod-to-Rod Bipolar Synapse. Neuron 41, 431–443.

Schneeweis, D.M., Schnapf, J.L., 1999. The photovoltage of macaque cone photoreceptors: adaptation, noise, and kinetics. J Neurosci 19, 1203–1216.

Schreyer, H.M., Gollisch, T., 2021. Nonlinear spatial integration in retinal bipolar cells shapes the encoding of artificial and natural stimuli. Neuron 109, 1692–1706.e8.

Shadlen, M.N., Kiani, R., 2013. Decision making as a window on cognition. Neuron 80, 791–806. 10.1016/j.neuron.2013.10.047

Turner, M.H., Rieke, F., 2016. Synaptic Rectification Controls Nonlinear Spatial Integration of Natural Visual Inputs. Neuron 90, 1257–1271. 10.1016/j.neuron.2016.05.006

Van Der Linde, I., Rajashekar, U., Bovik, A.C., Cormack, L.K., 2009. DOVES: a database of visual eye movements. Spat Vis 22, 161–177.

Verweij, J., Dacey, D.M., Peterson, B.B., Buck, S.L., 1999. Sensitivity and dynamics of rod signals in H1 horizontal cells of the macaque monkey retina. Vis. Res 39, 3662–3672.

Yu, Z., Turner, M.H., Baudin, J., Rieke, F., 2022. Adaptation in cone photoreceptors contributes to an unexpected insensitivity of primate On parasol retinal ganglion cells to spatial structure in natural images. eLife 11, e70611. 10.7554/eLife.70611

